# Mapping critical habitat of waterbirds in the Arctic for risk management in respect of IFC PS6

**DOI:** 10.1101/206763

**Authors:** Lammert Hilarides, Tom Langendoen, Stephan Flink, Merijn van Leeuwen, Bart Steen, Alexander V. Kondratyev, Andrea Kölzsch, Tomas Aarvak, Helmut Kruckenberg, Didier Vangeluwe, Emil Todorov, Anne Harrison, Eileen Rees, Adriaan M. Dokter, Bart Nolet, Taej Mundkur

**Affiliations:** Wetlands International, P.O. Box 4716700 AL Wageningen, The Netherlands; Affiliation Kondratyev; Max Planck Institute for Ornithology, Radolfzell, Germany; University of Konstanz, Konstanz, Germany; Norwegian Ornithological Society; Institute for Wetlands and Waterbird Research (IWWR) e.V., Germany, Am Steigbügel 3, D-27283 Verden (Aller), Germany

**Author notes:** ADDRESSES.

**Keywords:** Species distribution models, International Finance Corporation, Performance standards, Risk assessment

## Abstract

1. Economic development and energy exploration are increasing in the Arctic. Important breeding habitats for many waterbird species, which have previously been relatively undisturbed, are now being subjected to these anthropogenic pressures. The conservation of the habitats and the species they support is a significant challenge for sustainable development. Even if governments and corporates operating in this fragile environment are committed to sustainable development, there is little information available to avoid, mitigate and manage environmental risk and impacts. Taking a risk management perspective, we followed the International Finance Corporations’ (IFC) Performance Standard 6 (PS6) criteria on Environmental and Social Sustainability and developed an approach to identify “critical habitat”, as defined in IFC PS6, for waterbird species breeding in the Arctic. While the range of these waterbirds is roughly known, more accuracy is needed for proper risk assessment.
2. We have therefore gone a step further by modelling suitable habitat within these ranges. Depending on the relevance of the species for IFC PS6 and the level of certainty we separated the classes *likely* and *potential* critical habitat. We tested the approach for Russian breeding populations of five Anatidae species (White-fronted Goose *Anser albifrons*, Lesser White-fronted Goose *Anser erythropus*, Brent Goose *Branta bernicla*, Redbreasted Goose *Branta ruficollis* and Bewick’s Swan *Cygnus columbianus bewicki*). *Likely* critical habitats were identified through a review of literature and available data for these waterbird species and multi-species congregations. To address the information gap for most of the Russian Arctic a species distribution modelling approach was used. The outputs of this approach were labelled as *potential* critical habitat, indicating the lower level of certainty than *likely* critical habitat.
3. Based on existing information the amount of likely critical habitat is estimated to be at least x,xxx,xxx km^2^. For the five Anatidae species, X,XXX,XXX km^2^ potential critical habitat was identified; 95% of these areas were outside of the area boundaries of likely critical habitat for the species.
4. Insufficient data in the east of the study area did affect the results, as some areas known to support breeding populations were not identified as suitable. Conversely, species’ distributions may be overpredicted in other areas; It should also be recognized that the analyzed species currently have depressed populations and may therefore only utilize a proportion of suitable habitat available.
5. For risk assessment purposes however, it is better to predict false positives, rather than false negatives. The study indicates that there are large areas in the Arctic that are potentially important for each of the Anatidae species modelled, but are not yet recognised as key important areas. The results confirm that there is still much to learn about waterbird distribution and abundance in the Russian Arctic.
6. *Synthesis and applications* The critical habitat maps produced do not just provide a new source of information for the economic development sector, but provide it in a way that is relevant to the sector and directly applicable. The maps are useful for initial risk assessments of potential developments, to identify likely impacts and to consider mitigation options, in accordance with IFC PS6. Risk assessors should exercise caution and detailed surveys for any development in areas predicted to be suitable for each species should be carried out.

## INTRODUCTION

The Arctic provides important breeding habitat for many waterbird species that occur in Europe and Africa (Wohl 2006). Until recently, the breeding habitats have been relatively undisturbed, with low human densities, especially in comparison to other parts of the waterbird species’ flyways, where they compete with humans and many of their habitats have been modified or lost. However, with economic development, and oil and gas exploration, the Arctic is being subjected to increasing anthropogenic pressures that pose significant challenges for the management and conservation of Arctic habitats and the species they support (Wohl 2006).

The International Finance Corporation (IFC) is a member of the World Bank Group that focuses on private sector development and has a strategic commitment to sustainable development. For this, the IFC has developed eight performance standards on social and environmental sustainability; approximately 80 large corporates in the primary resource and financing sectors have adopted these standards. Of relevance to the protection of habitats and waterbirds in the Arctic, Performance Standard 6 (PS6) deals with “Biodiversity conservation and sustainable management of living natural resources” (IFC 2012a). In accordance with IFC PS6 different risk management approaches are employed to protect biodiversity and ecosystem services based on the sensitivity and values of a habitat. Thus, the identification of important habitats for waterbirds is a crucial step to inform management plans and minimise the impacts of human activities.

PS6 gives a definition of “critical habitat” and provides guidance on how to act when operating in or close to a critical habitat (IFC 2012b). Critical habitat is a geographic area important for biodiversity and may include: (1) habitats of significant importance to Critically Endangered and/or Endangered species (as categorized in the IUCN Red List of Threatened Species; IUCN 2015); (2) habitats of significant importance to endemic and/or restricted range species; (3) habitats that support globally significant concentrations of migratory species and/or congregatory species; (4) highly threatened and/or unique ecosystems; and/or (5) areas associated with key evolutionary processes (IFC 2012a). If an area contains critical habitat, IFC PS6 requires a Biodiversity Action Plan to be developed and implemented (IFC 2012a). However, business developers rely on existing species distribution, biodiversity and protected areas data sets, because there is no global map of critical habitat. While some of the existing data sets are good indicators of critical habitat and use criteria that overlap with those used by the IFC (Martin *et al.* 2015), the data are generally incomplete or require interpretation under the IFC guidelines. As a result, there are many areas of critical habitat for species, ecosystems and evolutionary processes that have not yet been identified, particularly in the Arctic. Only few robust, long-term monitoring programmes are in action or openly available here, even for waterbirds, one of the most intensely studied animal groups in the world.

In this study, we propose a new methodology to identify critical waterbird habitat in the Arctic, based on PS6 criteria. We focused on areas covered by both the Conservation of Arctic Flora and Fauna (CAFF) working group and the African Eurasian Waterbird Agreement (AEWA). Given the limited data availability and geographic gaps in information, we adopted a modelling approach. The model outputs are translated into maps that detail potential and likely areas of IFC PS6 critical habitat in the Arctic. These maps can improve conservation by supporting risk assessments for potential developments, identify likely impacts and consider mitigation options.

## MATERIALS AND METHODS

### Species selection

We analysed five Anatidae species: White-fronted Goose *Anser albifrons*, Lesser White-fronted Goose *Anser erythropus*, Brent Goose *Branta bernicla*, Redbreasted Goose *Branta ruficollis* and Bewick’s Swan *Cygnus columbianus bewicki*. These all have populations that breed exclusively in the Russian Arctic and were considered likely to trigger criteria 1 or 3 of PS6, based on their red list status or occurrence in large enough concentrations, respectively.

### Model

To produce detailed species distribution maps that could serve as a basis for critical habitat maps we used MaxEnt (Phillips, Dudík & Schapire 2004). The maps output by the model predict the suitability of habitat in the study area. The model has been used widely in the scientific community for a variety of species across a wide spectrum of habitats (Elith *et al.* 2006; Phillips & Dudik 2008;Edrén *et al.* 2010).

### Species occurrence

Species occurrence samples were obtained from online databases, telemetric studies, regional surveys and literature sources. To exclude data on vagrant birds, or otherwise unrepresentative data, only samples from within the known range of a species were used. The large distances migratory waterbirds cover during migration means that they use very different habitats during different life cycle stages, including breeding, moulting, migration and wintering. Combining these life cycle stages into a single model would lead to an overprediction of suitable habitats. The data were therefore filtered by date and location, specific for each species, so that only the breeding season samples remained (see Fig. S1 in Supporting Information). Breeding season samples may include moulting for some species, when moulting happens at the same location. As such, samples were principally categorized as breeding or breeding/moulting.

Especially when using samples from online databases, such as the Global Biodiversity Information Facility (GBIF), filtering the data (as described above) is an essential step. To illustrate this, none of the 448 occurrence samples of *B. ruficollis* in the GBIF from June, July or August fall within the known breeding range of the species and are likely observations of escaped captive birds and vagrants. To address IFC PS6 Criterion 3, it was also considered important to distinguish whether a species congregates during a particular life cycle stage, as is often the case during migration and moulting, and sometimes also during breeding.

**Telemetry data**: while telemetry data provide occurrence samples that represent true and relatively accurate occurrences of individuals, these data may not be representative of the full population because of the limited number of individuals equipped with transmitters. By design, these samples are also highly autocorrelated; that is, every successive sample is inherently close to the preceding sample, both in time and space. Since MaxEnt assumes a random distribution of occurrence samples, this affects the model quality (Phillips *et al.* 2009). As demonstrated by (Fourcade *et al.* 2014), applying a spatial filter is a relatively good, and is the most consistently performing, method to mitigate the effects of sample bias. In addition, the use of multiple data sources mitigated the effects of the potentially unrepresentative telemetry-based samples.

### Environmental predictors

Environmental predictors were selected that would potentially influence the species’ distributions, were available and were of consistent quality across the entire study area. Selected environmental predictors included bio-climatic variables, distances to different types of waterbodies, elevation, soil-related variables and land cover data (see Table S1 in Supporting Information). To prevent distortion of the model by decreasing raster cell size at higher latitudes (Elith *et al.* 2011), all predictors were harmonised in a GIS by re-projecting to an equal area projection (the North Pole Lambert Azimuthal Equal Area; EPSG: 102017) at a 1 km resolution.

Although many environmental predictors were readily available from the literature or online databases, some environmental predictors, expected to be of ecological significance to each of the species, were produced (distances to coast, freshwater, estuary and shallow coastal flats, slope, terrain roughness and dominant soil type; see below). Because of the different sources of data, the exact extent (generally the coastline), of each produced environmental predictor was not consistent, so the extent of the bio-climatic variables was used as a reference.

**Distance to coast:** This predictor was created by measuring the “Euclidian distance” to sea, using a bioclim predictor as reference for the coastline.

**Distance to freshwater:** this was based on the 250 m MODIS Water Mask data set (Carroll *et al.* 2009) with the sea masked out using the “no data” zone of a bioclim predictor. The “Euclidian distance” tool was used to calculate the distance between each cell in the study area and the nearest cell with freshwater.

**Distance to estuary:** although there is a Global Estuary Database (Alder 2003), this was considered too coarse, with many medium to small estuaries omitted. Therefore, the lowest sub-basin polygon was selected from the HydroBASINS level 10 data set (Lehner & Grill 2013), for each basin larger than 500 km^2^. This minimum basin size threshold was set to avoid the selection of every coastal polygon. The “Euclidian distance” was then calculated between each cell in the study area and the nearest estuary.

**Distance to shallow coastal flats:** shallow coastal flats function as feeding grounds for many waders, (sea-) ducks and geese. The International Bathymetric Chart of the Arctic Ocean (Jakobsson *et al.* 2012) was used and the shallow areas at sea (experimentally determined between +1 m and −1 m) were selected. Within these shallow areas a sub-selection was made of all the areas that had a slope less than 0,002. From this subset only areas of at least 10 km^2^ were selected. The “Euclidian distance” tool was used to calculate the distance between each cell in the study area and the nearest shallow coastal flat.

**Slope:** this was calculated from the BIOCLIM digital elevation model.

**Terrain roughness:** the standard deviation predictor of the 30 arc-seconds USGS Global Multiresolution Terrain Elevation Data (GMTED2010) (Danielson & Gesch 2011) was used as a proxy for terrain roughness. A “rough” area has large fluctuations in height and therefore a large standard deviation of elevation, whereas flat areas have a low standard deviation.

**Dominant Soil Type:** this predictor was based on the International Soil Reference and Information Centre’s 1 km soilgrid data set (Hengl *et al.* 2014). Unfortunately, this predictor had data gaps for areas covered with seawater and freshwater, as well as permanent snow and ice cover. These gaps were partially filled using freshwater pixels from 250m MODIS Water Mask.

#### Correlation analysis

Although MaxEnt is relatively robust to correlated predictors (Elith *et al.* 2011), removing them does tend to improve the model (Elith & Leathwick 2009). Therefore, R was used to conduct pairwise assessments of the correlations between the environmental predictors (Table S2). Because not all predictors had a normal distribution, Spearman’s rank correlation coefficients between each possible combination of predictors based on 100,000 random locations were calculated.

### Test model runs

Test models were run using the quality assessed and spatially filtered occurrence samples and all environmental predictors, to identify the most important predictors for each species model (Table S1). Predictors were ranked by their permutation importance, as an indication of unique information, and percent contribution. The highly correlated predictors, with Spearman rho values below −0.75 or above 0.75 were removed (Table S2); this meant the predictors with the highest permutation importance were retained.

### Model runs

MaxEnt was run with the selected environmental predictors and the following settings changed from their default: (1) 10 replicates; (2) bootstrap sampling; (3) random seed; (4) 30% random test percentage; (5) response curves; (6) jackknife procedure; (7) maximum iterations were set at 2000 and 8) write background predictions.

### Suitable habitat

One output of MaxEnt for each model run was a map, showing the average probability (of the 10 replicates) of habitat suitability for each species within each raster cell; that is, a logistic value between 0 (not suitable) and 1 (very suitable) (Fig. S1). A threshold was applied to these probability maps for each species to create binary suitable/unsuitable habitat maps. The threshold was calculated using the “equal training sensitivity and specificity” method in MaxEnt, to provide a balance between the omission and commission errors.

### Critical habitat

This study applied the criteria from the IFC PS6 Guidance Notes using a rules based approach (Table 1) to classify critical habitat from the maps showing suitable habitats. These rules were derived from IFC PS6 criteria 1 and 3. In areas other than the Arctic, or for species groups other than waterbirds, criterion 2 (on endemic or restricted range species) might also be applicable.

**Table 1.**
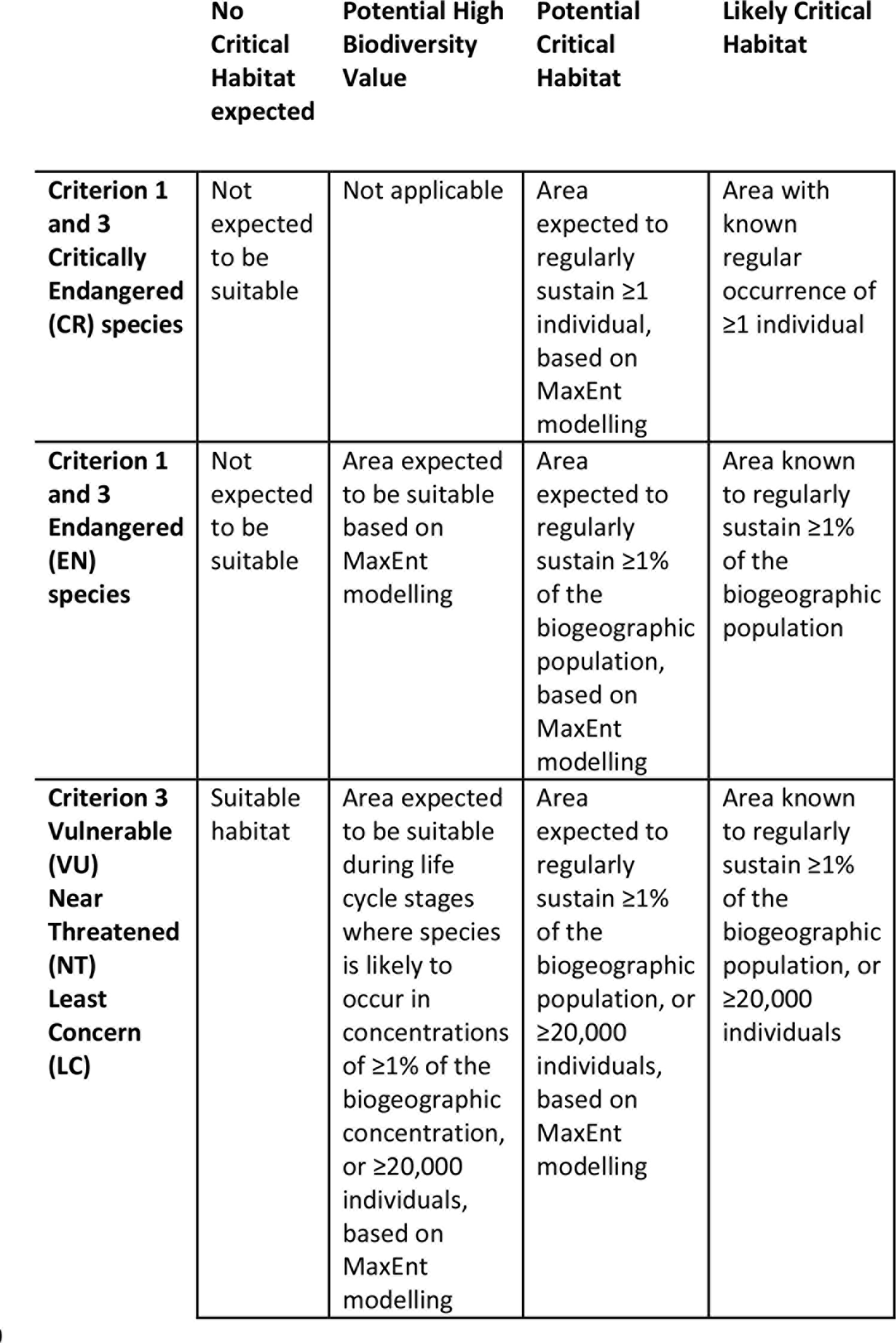
Classification and justification of critical habitat based on IFC PS6

|  | No<br>Critical<br>Habitat<br>expected | Potential High<br>Biodiversity<br>Value | Potential<br>Critical<br>Habitat | Likely Critical<br>Habitat |
| --- | --- | --- | --- | --- |
| **Criterion 1**<br>**and 3**<br>**Critically**<br>**Endangered**<br>**(CR) species** | Not<br>expected<br>to be<br>suitable | Not applicable | Area<br>expected to<br>regularly<br>sustain $\geq 1$<br>individual,<br>based on<br>MaxEnt<br>modelling | Area with<br>known<br>regular<br>occurrence of<br>$\geq 1$ individual |
| **Criterion 1**<br>**and 3**<br>**Endangered**<br>**(EN)**<br>**species** | Not<br>expected<br>to be<br>suitable | Area expected<br>to be suitable<br>based on<br>MaxEnt<br>modelling | Area<br>expected to<br>regularly<br>sustain $\geq 1\%$<br>of the<br>biogeographic<br>population,<br>based on<br>MaxEnt<br>modelling | Area known<br>to regularly<br>sustain $\geq 1\%$<br>of the<br>biogeographic<br>population |
| **Criterion 3**<br>**Vulnerable**<br>**(VU)**<br>**Near**<br>**Threatened**<br>**(NT)**<br>**Least**<br>**Concern**<br>**(LC)** | Suitable<br>habitat | Area expected<br>to be suitable<br>during life<br>cycle stages<br>where species<br>is likely to<br>occur in<br>concentrations<br>of $\geq 1\%$ of the<br>biogeographic<br>concentration,<br>or $\geq 20,000$<br>individuals,<br>based on<br>MaxEnt<br>modelling | Area<br>expected to<br>regularly<br>sustain $\geq 1\%$<br>of the<br>biogeographic<br>population, or<br>$\geq 20,000$<br>individuals,<br>based on<br>MaxEnt<br>modelling | Area known<br>to regularly<br>sustain $\geq 1\%$<br>of the<br>biogeographic<br>population, or<br>$\geq 20,000$<br>individuals |

The classification for critical habitat followed that of Martin (Martin *et al.* 2015) and distinguished between “potential” and “likely” critical habitat, based on relevance and certainty, indicating the difference between *modelled* critical habitat and that *confirmed* by literature or other sources.

**Likely critical habitat:** Independently from the modelling study, for each species and life cycle stage, areas that would qualify as critical habitat under IFC PS6 were identified through a review of the literature (Krivenko 2000);(Ramsar Convention);(Arctic and Antarctic Research Institute), from IBA information (Birdlife International) and from the Critical Site Network Tool (Wings Over Wetlands 2011). Under PS6 criteria an area qualifies as critical habitat if it regularly supports 1% of a migratory species’ population. The areas known to support >1% of a population were therefore classified as “likely critical habitat” and also served as input data for the next step, in which the threshold value for “potential critical habitat” was determined.

**Potential critical habitat:** The threshold value used to identify the “suitable habitat” for a species (see above) was based on statistical, rather than ecological, considerations. The “suitable habitat” had quite a large range of probability values; that is, from 0.16 for the “least probable suitable habitat” to 0.92 for the “most probable suitable habitat”. To parameterize IFC PS6 criterion 3 and identify “potential critical habitat”, the habitat that was suitable enough as well as large enough to support congregations of >1% of a species’ population was identified; that is, habitat that was (1) more suitable than the average habitat suitability of known key areas; and (2) larger than the typical size of known key areas was identified. For this, the key areas identified as likely critical habitat were overlaid with the probability map and then: (1) the average probability value of the suitable habitat within the key areas was calculated; and (2) the median size of the suitable habitat within the key areas was calculated. For (2), the median was preferred over the average, because of the small sample size and to minimize the effect of extreme values. The raster was then resampled from the original 1-km resolution grid, using the square root of the median of the suitable habitat area in the known key areas, to identify suitable habitat of sufficient size. The average probability value calculated in step (1) was then used as the threshold to identify the potential critical habitat on the resampled raster. Thus, the resulting maps identified habitats with a relatively high probability of meeting PS6 criteria for critical habitat, based on their suitability and size.

### Validation

As demonstrated by Termansen (Termansen, McClean & Preston 2006) and Lobo (Lobo, Jiménez-valverde & Real 2008), measuring model accuracy solely from the often used “Area Under Curve” (AUC) values may be misleading. Therefore, to further validate our results, we used the “True Skill Statistic” (TSS) (Allouche, Tsoar & Kadmon 2006) and additionally also list the sensitivity and specificity. These scores are based on the 30% random test percentage of the species occurrence samples. In addition the maps were validated during expert reviews conducted by the relevant Wetlands International expert groups.

## RESULTS

### Overview

A total of xxx species occurrence samples were collected and, after spatial filtering, 740 were used, with a minimum of 90 samples per species model (Table 2; Fig. 1). There were a limited number of samples in the east of the study site. The AUC score for all models was >0.9 and, importantly, the sensitivity, specificity and TSS scores were also good to very good (Table 3). A total of 1,767,749 km^2^ within the study area was identified as potential critical habitat from a total of 4,599,512 km^2^ identified as suitable habitat (Tables 1 & 4). 95% of the potential critical habitat and 96% of the other areas of suitable habitat were outside the boundaries of areas *known* to hold 1% of the species (Table 4; Figs 2 & 3).

**Table 2.** Occurrence samples of each species used

| Species | # occurrence samples | 1% area |
| --- | --- | --- |
| *Anser albifrons*<br>(Greater White-fronted Goose) | 176 | 1,672 |
| *Anser erythropus*<br>(Lesser White-fronted Goose) | 90 | 499 |
| *Branta bernicla*<br>(Brent Goose) | 116 | 390 |
| *Branta ruficollis*<br>(Red-breasted Goose) | 210 | 749 |
| *Cygnus columbianus*<br>(Tundra Swan) | 148 | 623 |

**Table 2a.**
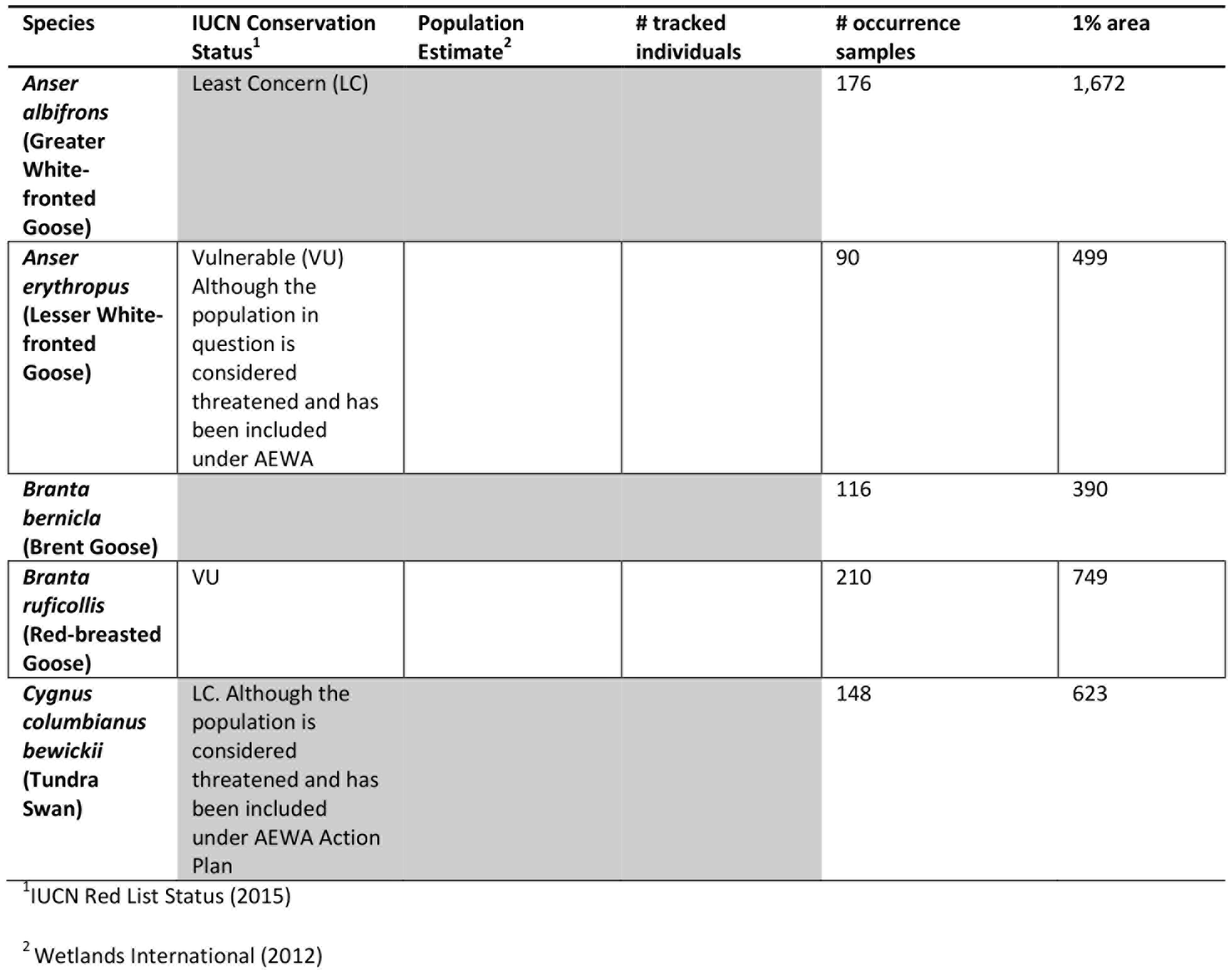
Overview of species and occurrence samples used

**Fig. 1.**
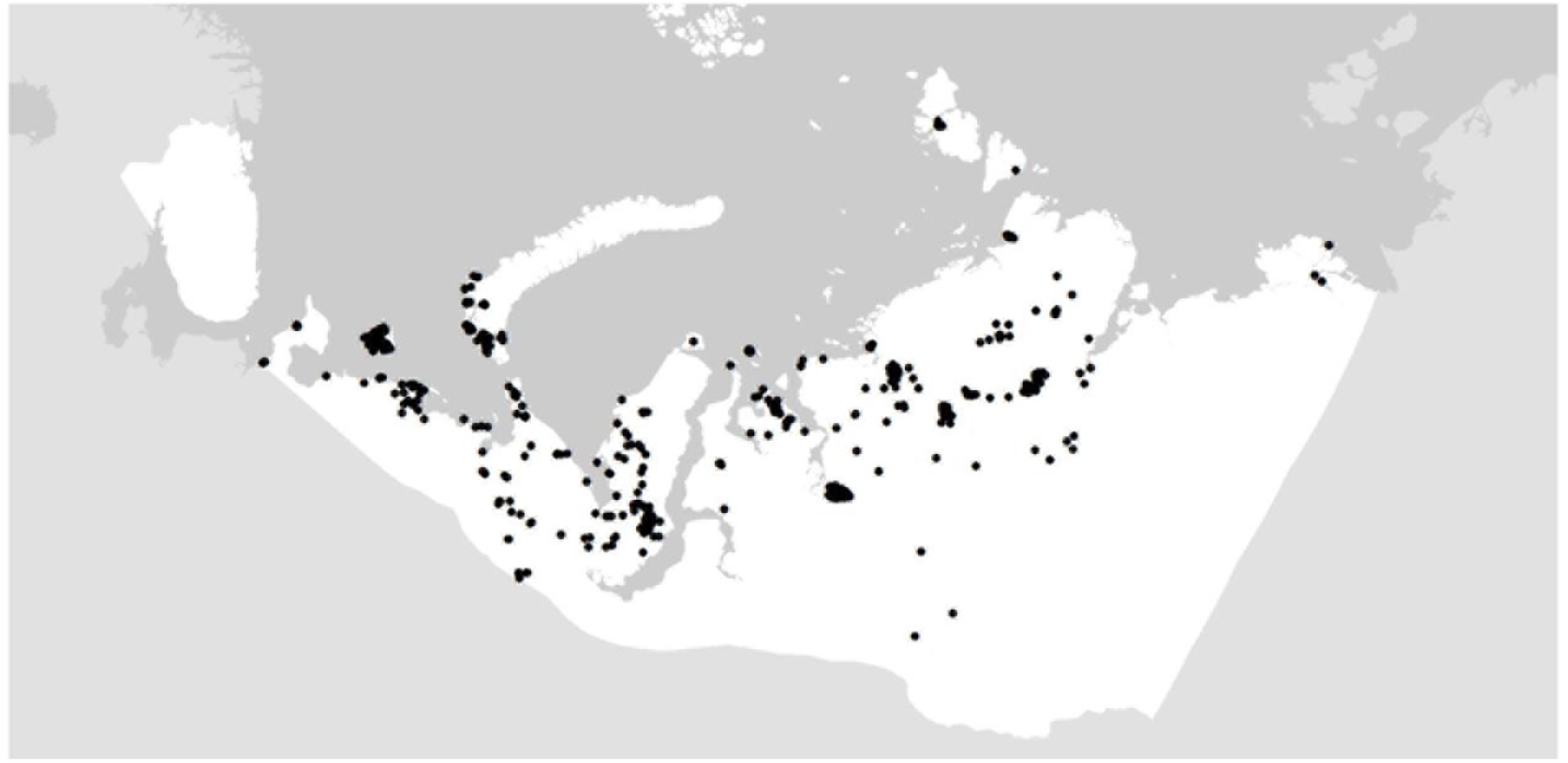
Distribution of waterbird occurrence samples used (black dots)

**Table 3.**
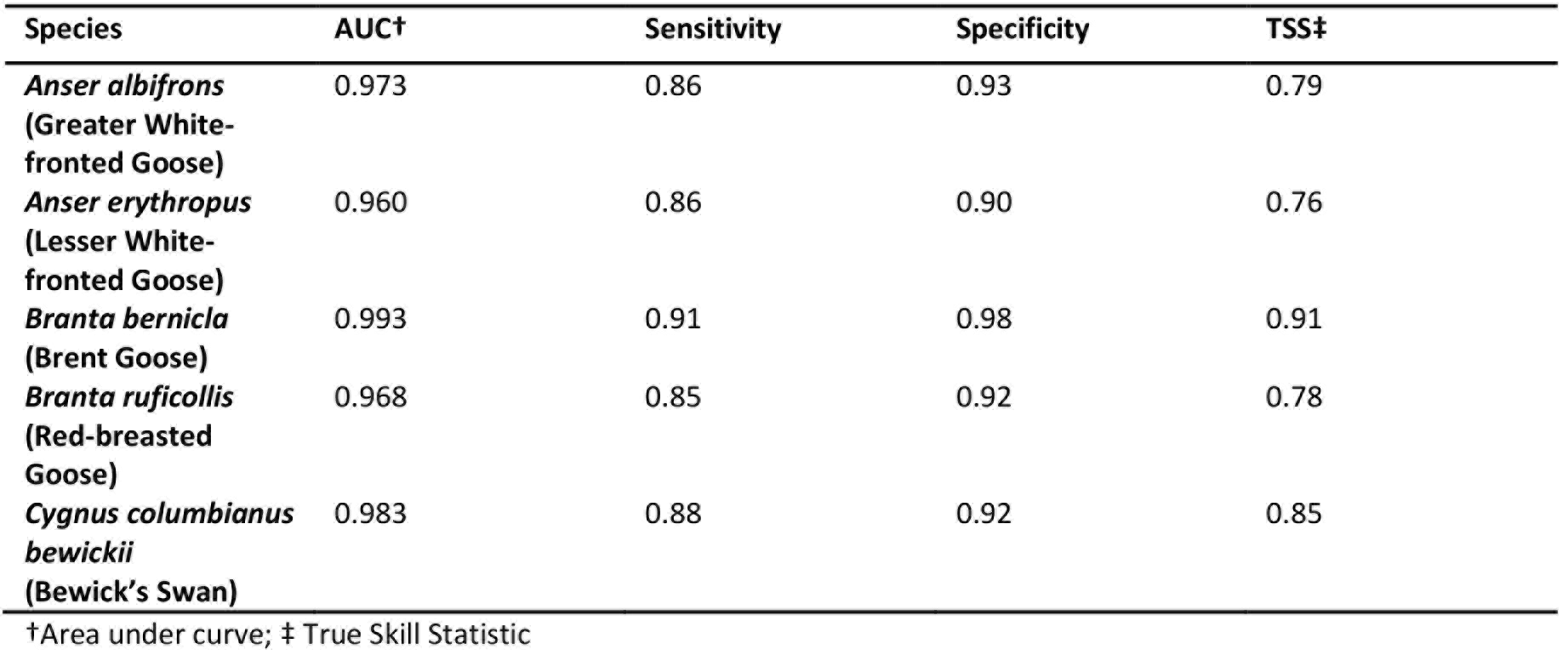
Validation of the MaxEnt models

| Species | AUC† | Sensitivity | Specificity | TSS‡ |
| --- | --- | --- | --- | --- |
| *Anser albifrons* (Greater White-fronted Goose) | 0.973 | 0.86 | 0.93 | 0.79 |
| *Anser erythropus* (Lesser White-fronted Goose) | 0.960 | 0.86 | 0.90 | 0.76 |
| *Branta bernicla* (Brent Goose) | 0.993 | 0.91 | 0.98 | 0.91 |
| *Branta ruficollis* (Red-breasted Goose) | 0.968 | 0.85 | 0.92 | 0.78 |
| *Cygnus columbianus bewickii* (Bewick's Swan) | 0.983 | 0.88 | 0.92 | 0.85 |

†Area under curve; ‡ True Skill Statistic

**Table 4.** Potential high biodiversity value (HBV) and potential critical habitat (PCH) per species, and the fractions in known important areas (KIA)

| Species | # KIA | PCH (km<sup>2</sup>) | HBV (km<sup>2</sup>) | PCH outside<br>KIA km<sup>2</sup> (%) | HBV outside<br>KIA km<sup>2</sup> (%) | KIA with<br>PCH # (%) | KIA with<br>HBV # (%) |
| --- | --- | --- | --- | --- | --- | --- | --- |
| Anser albifrons<br>(Greater<br>White-fronted<br>Goose) | 15 | 164,635 | 1,235,684 | 144,478<br>(88%) | 1,156,328<br>(94%) | 4<br>(27%) | 13<br>(87%) |
| Anser<br>erythropus<br>(Lesser White-fronted Goose) | 11 | 457,225 | 916,111 | 445,888<br>(98%) | 893,747<br>(98%) | 5<br>(45%) | 8<br>(73%) |
| Branta bernicla<br>(Brent Goose) | 10 | 223,979 | 385,941 | 221,811<br>(99%) | 377,015<br>(98%) | 2<br>(20%) | 5<br>(50%) |
| Branta<br>ruficollis<br>(Red-breasted<br>Goose) | 13 | 533,249 | 1,162,723 | 516,439<br>(97%) | 1,123,013<br>(97%) | 4<br>(31%) | 9<br>(69%) |
| Cygnus<br>columbianus<br>(Tundra Swan) | 11 | 135,217 | 958,341 | 115,843<br>(86%) | 908,903<br>(95%) | 3<br>(27%) | 10<br>(91%) |
| Total distinct<br>area |  | **1,767,749** | **4,599,512** | **1,674,434**<br>**(95%)** | **4,419,112**<br>**(96%)** |  |  |

**Fig. 2.**
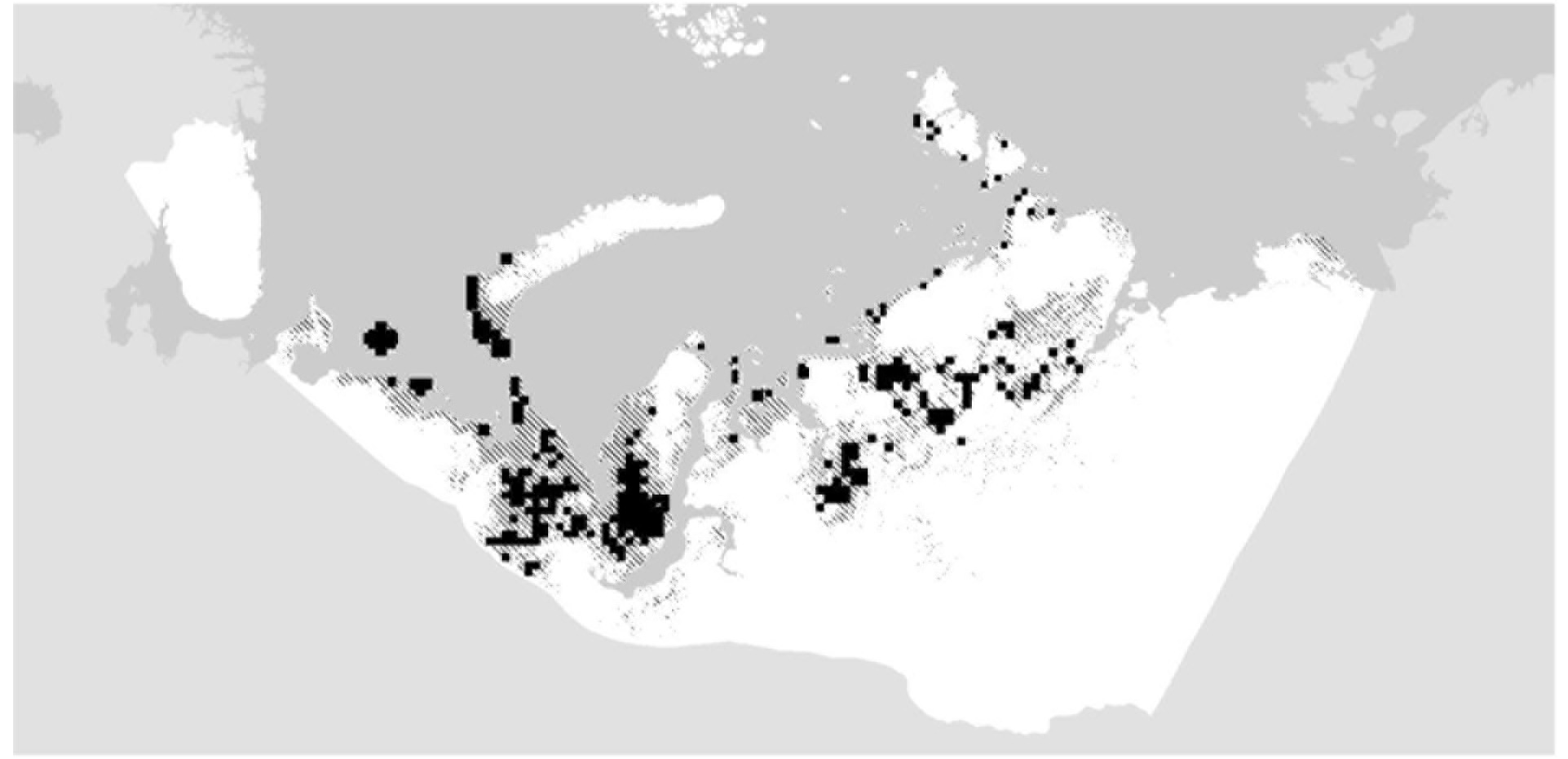
Suitable habitat (dashed) and potential critical habitat (black) during the breeding season for all five species

**Fig. 3.** Known important areas for all species

### Anser albifrons

As one of the most common geese in the Russian arctic, *A. albifrons* had a high number of occurrence records across most of its known range (Fig. 4a), although fewer occurrence points were available for the populations that occur east of the Taimyr. The model predicted suitable breeding habitat for large concentrations across much of the known breeding range (Wings Over Wetlands Project 2010) and 13 of the 15 known important areas for the species (Fig. 4b). The two areas that were not predicted to have suitable breeding habitat were in the far east of the study area. A total of 21 areas of potential critical habitat were identified, of which 11 overlapped with known critical habitat. Many of the newly identified areas were also between or adjacent to known critical habitat areas, such as along the coast of Baydaratskaya Guba, although 3 new areas were identified in Nova Zemblya.

**Fig. 4.**
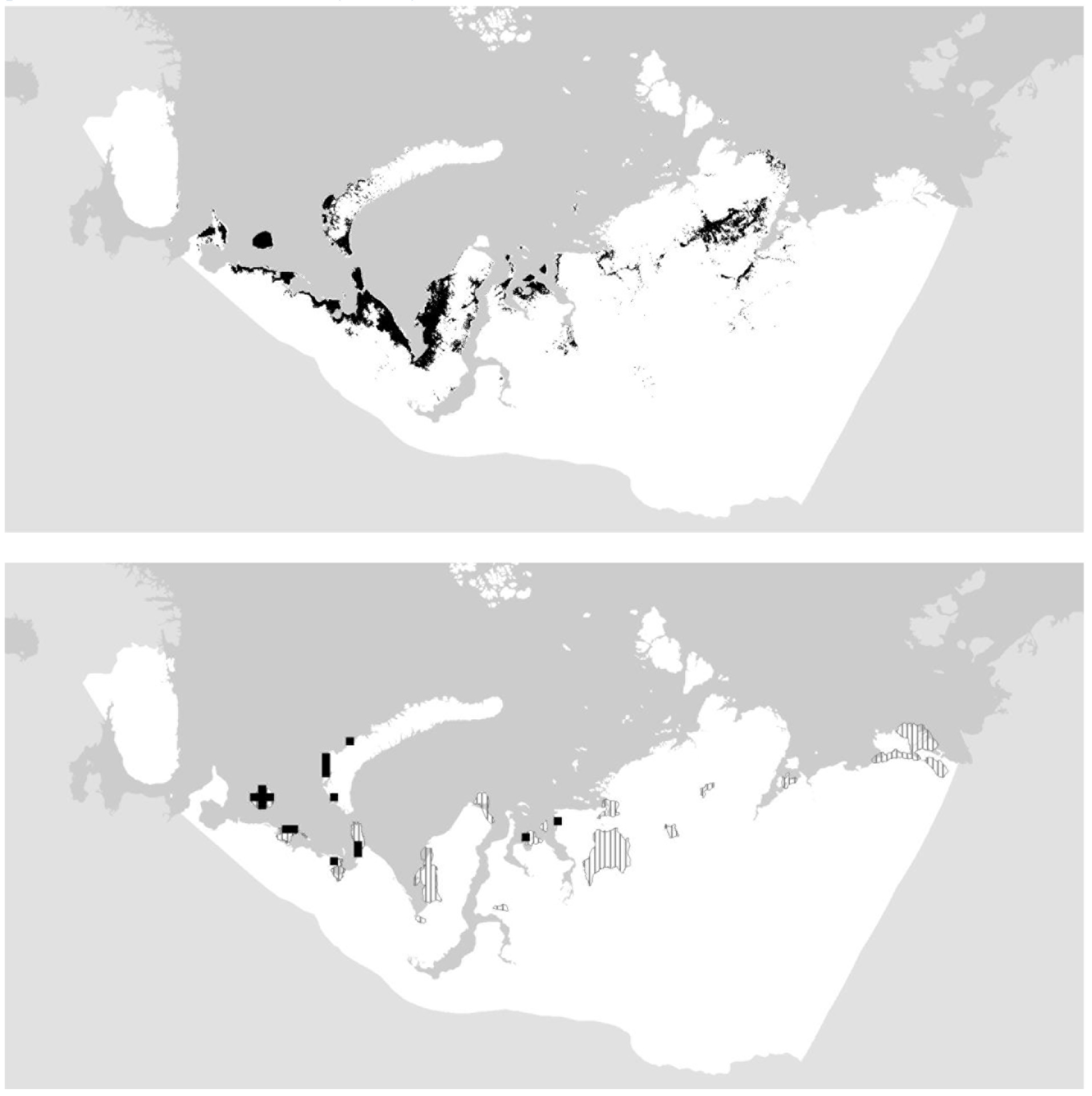
*Anser albifrons* (a) identified suitable habitat, and (b) likely (dashed) and potential critical habitat (black)

### Anser erythropus

*A. erythropus* had the fewest number of occurrence records (90) in the study. No occurrence points were available for the most easterly population known, which is located in Central and Eastern Siberia (Fig. 5a). Additionally, the populations of *A. erythropus* that breed in Russia have declined rapidly (BirdLife 2015 XX1), which may affect the representativeness of the historical samples. Large areas outside of the known breeding range (Wings Over Wetlands Project 2010) were predicted as potential critical habitat, especially in the Yamal and Yugorskiy Peninsulas (Fig. 5b). Conversely, only one known key area in the Taimyr was identified as potential critical habitat, although the area predicted as suitable in the Taimyr closely matched the known breeding range.

**Fig. 5.**
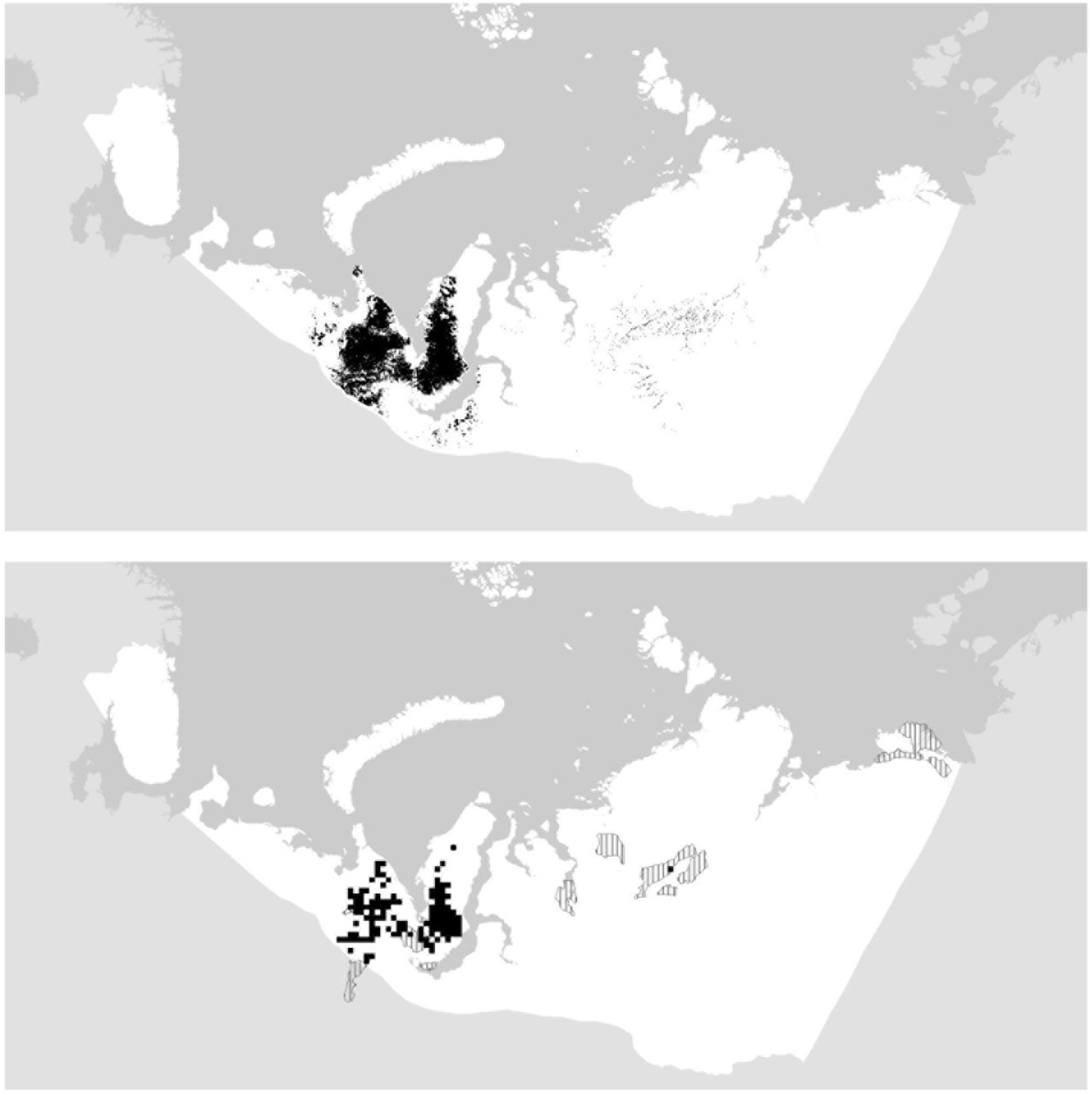
*Anser erythropus*(a) identified suitable habitat, and (b) likely (dashed) and potential critical habitat (black)

### Branta bernicla

This species breeds and moults close to the coast in the arctic tundra or islands (BirdLife 2015 XX2). Fewer occurrence samples (116) were available for this species than most of the others studied, with most records spread across the central part of the species’ breeding range (Fig. 6a). Areas between the eastern Taimyr and Lena Delta, Nova Zemblya, and the far west of the study area were underrepresented. Consequently, four of the most westerly sites, identified as known key areas, were not predicted as suitable, although these areas were peripheral or outside of the breeding range maps of the species (Fig. 6b) (Wings Over Wetlands Project 2010). In addition, three known important areas in the eastern part of the study area were not identified as suitable. However, nine areas were identified as new potential critical habitat in the islands to the far north, including Bolshevik Island, and a number of areas along the West Taimyr coast, up to the Yamal Peninsula.

**Fig. 6.**
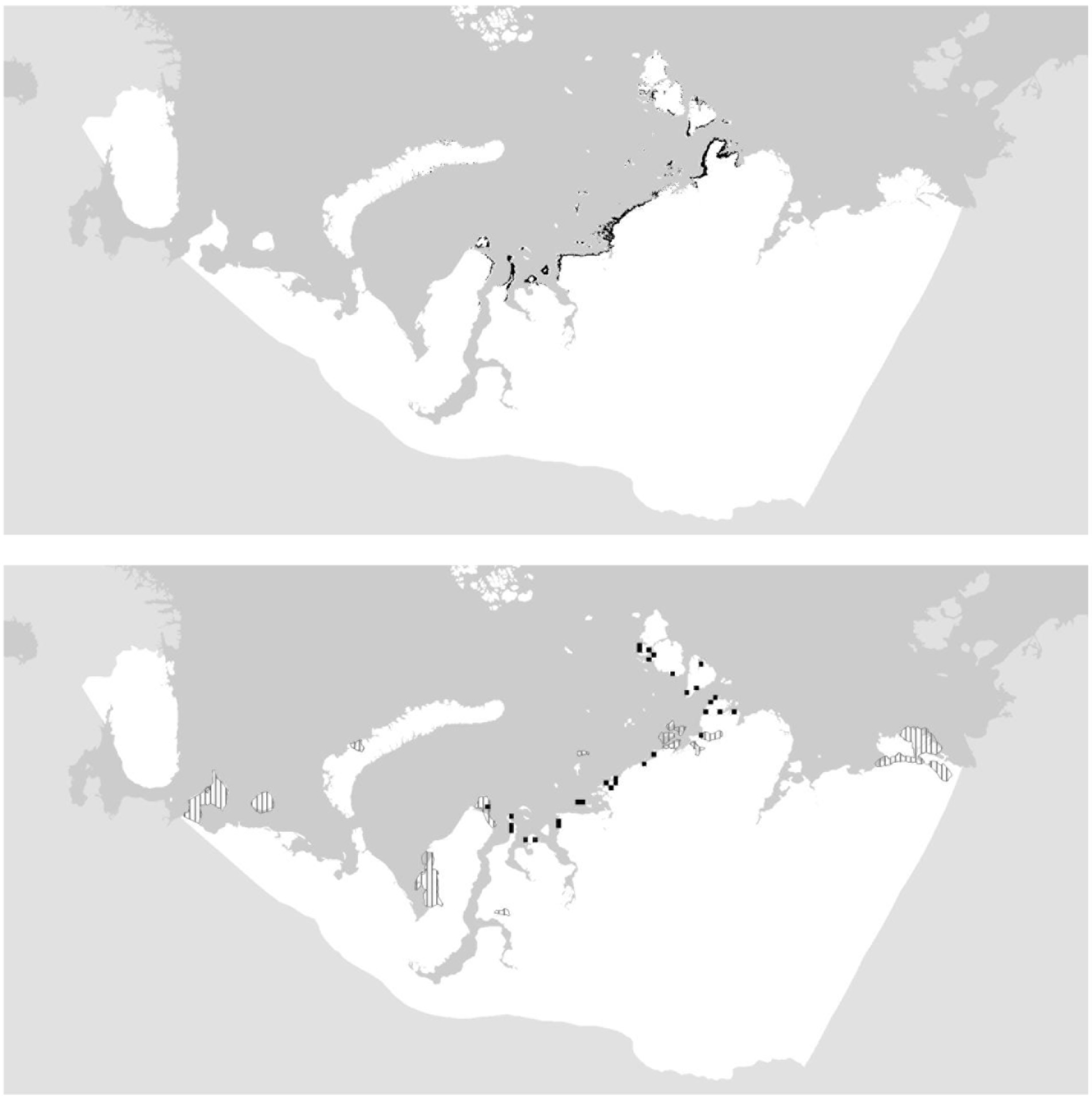
*Branta bernicla* (a) identified suitable habitat, and (b) likely (dashed) and potential critical habitat (black)

### Branta ruficollis

*B. ruficollis* has the smallest population size of the species studied (Table X) and its breeding range is restricted to areas in the Taimyr, Gydan and Yamal Peninsulas (BirdLife 2015 XX3). The species’ distribution was modelled from 210 occurrence samples (Fig. 7a), mostly from the Taimyr, where an estimated 70% of the population breeds (BirdLife 2015 XX3). In total, 11 known key areas were not identified as potential critical habitat by the species model (Fig. 7b); however, many of these areas were outside or on the very edge of the reported breeding range of the species (Wings Over Wetlands Project 2010). In addition, large new areas were predicted as potential critical habitat within the breeding range, mainly located in the Taimyr and parts of the Yamal Peninsula.

**Fig. 7.**
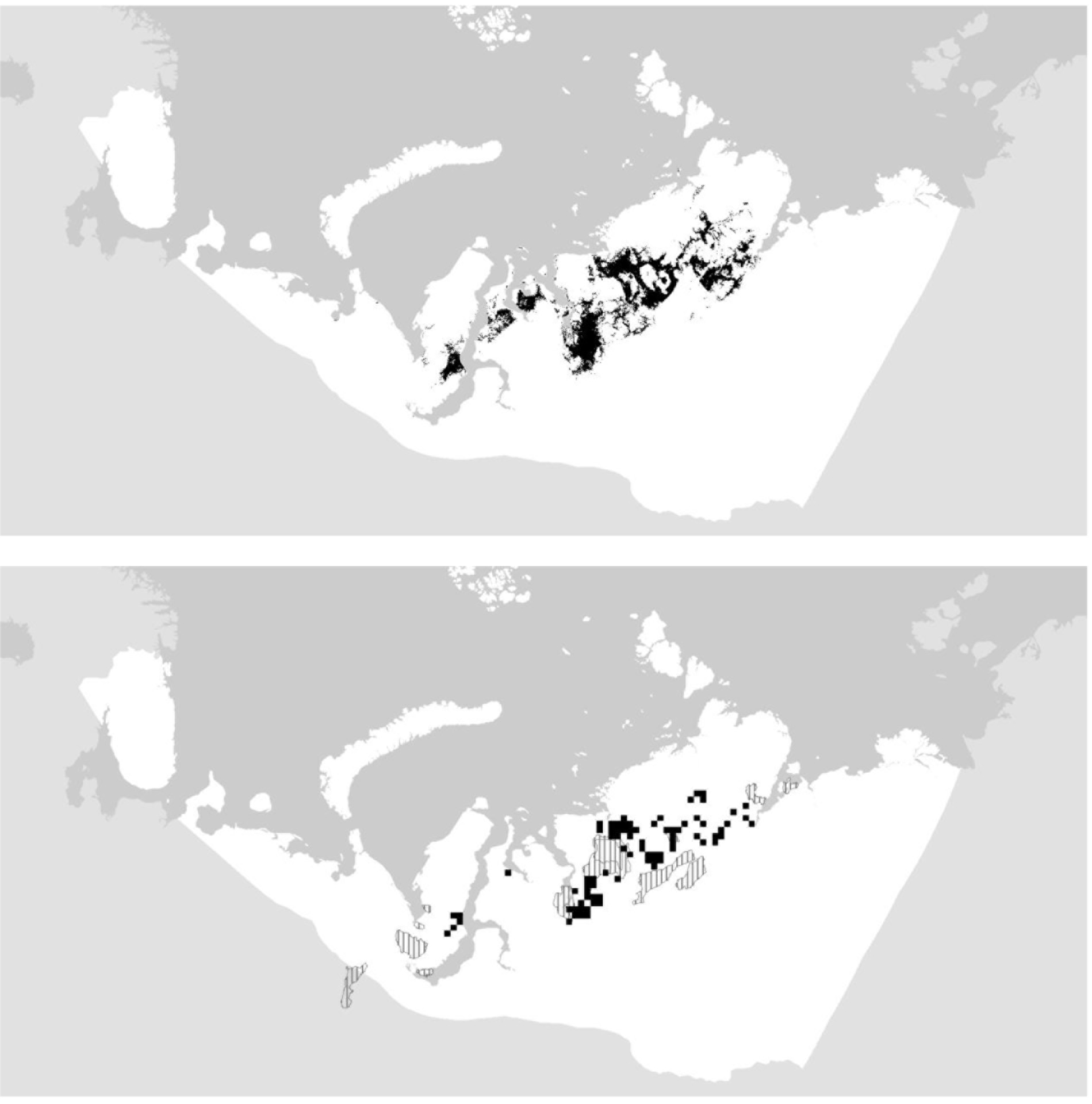
*Branta ruficollis* (a) identified suitable habitat, and (b) likely (dashed) and potential critical habitat (black)

### Cygnus columbianus bewickii

Occurrence samples for this species were predominantly from the Western Siberia and North-East/North-West Europe population, with far fewer samples from the Northern Siberia/Caspian and Asian populations (Wetlands International 2012), which breed in the Taimyr and to its east, up to the Lena Delta (Fig. 8a). Consequently, known key areas in the eastern portion of the study area were not identified as potential critical habitat, even though large parts of the Lena Delta were identified as suitable habitat (Fig. 8b). The known key areas in the west were identified as potential critical habitat, with the exception of areas in the south of the Yamal Peninsula, which is on the periphery of the Western Siberia and North-East/North-West Europe population range (Wings Over Wetlands Project 2010). New potential critical habitat was also identified in the southern part of Nova Zemblya, which is in accordance with the breeding range of the species (Wings Over Wetlands Project 2010)

**Fig. 8.**
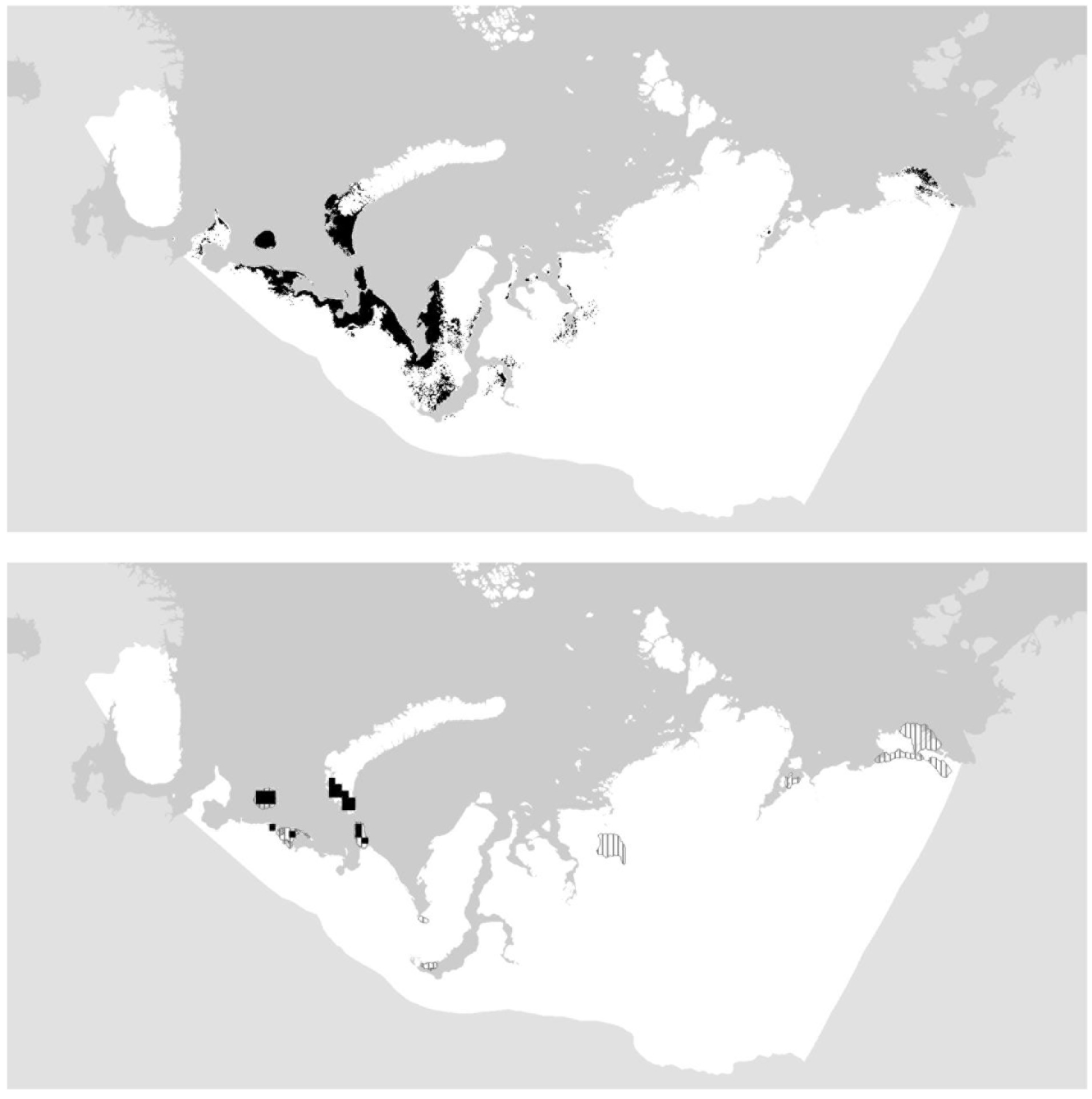
*Cygnus columbianus* (a) identified suitable habitat, and (b) likely (dashed) and potential critical habitat (black)

## DISCUSSION

Identifying suitable habitat through species using modeling methods is a long-standing and verified approach (Ref.) and can provide a powerful tool particularly in regions that are remote and data poor. We used the MaxEnt model to identify suitable habitats for congregations of five Anatidae species to be used for risk assessment purposes. The results are in broad agreement with the known breeding ranges of the species. About 95% of the potential critical habitat identified was outside known critical (?) areas, indicating that there are large areas in the Arctic which are potentially important for each of the species modelled. The current population size of the species may have a strong influence on the areas that are currently being favoured for breeding by a species, particularly in congregatory Anatidae. As a result, it may be expected that for those species with currently depressed populations (as compared to higher historic populations), the areas that are currently being utilised are considerably smaller than the overall suitable habitat available for a species in this part of the arctic. Therefore, it may seem there are some cases in our study where the species, distributions may appear to have been overpredicted. For example, *B. ruficollis* has a known population of only 55,000 (Wetlands International 2015). However the species is unlikely to occupy this entire area at the same time. Also, a key factor that determines *B. ruficollis* breeding areas is the presence of raptors (Prop & Quinn 2003); however, this environmental variable was not included in this study, which may have influenced the results. Nevertheless, for a risk assessment the results are valuable to identify areas where the species may be present, especially as *B. ruficollis* is listed as a Vulnerable species on the IUCN Red List. In addition, for risk assessments it is arguably better to predict false positives, rather than false negatives.

Another reason for the mismatch may be that known critical habitat areas need further verification.

Our study has only focused occurrence data from the breeding and moulting seasons of these species. Extending this approach to other crucial life cycle stages, particularly the pre-breeding (northward) and post breeding (southward) migration periods, when birds may congregate and require different habitats across the breadth of the arctic should provide an important basis for initial risk assessments of potential developments in these areas too.

The accuracy of occurrence points is vital for the model results, and as many of the occurrence records used were based on telemetry data or highly accurate survey techniques, this should enable overall positive results. However there were large areas that were not predicted as suitable within the overall known breeding ranges of the species. This could be a result of small population sizes of species, as mentioned above, differences in the habitat preferences across their range or insufficient data (occurrence samples) in the particular areas. The latter reason could particularly have affected species occupying the eastern part of the study area, where occurrence records were scarce.

Following the IFC PS6 Guidance Notes, we attempted to identify habitat that was not only suitable for congregations of waterbirds or endangered species, but also critical. Converting species modelled suitability into critical habitat has no precedence in the scientific literature and our approach can be considered highly conservative. By identifying new critical habitat using the median area and average probability of habitat identified as suitable in known critical habitats, we automatically set a threshold for potential critical habitat that would exclude half of the known critical habitat areas.

As a result, risk assessors should remain cautious of important areas that were not identified through the modelling process. We advise that any area predicted to be suitable for each species is surveyed in more detail, with particular attention to the areas predicted to be potentially critical.

While populations of bird species are known to vary over decades, nearly all five species are declining due to changes and pressures in the arctic and along their entire migration cycles. Furthermore, habitats across the arctic remains a highly dynamic state and are being greatly influenced by past and ongoing natural and human induced changes within the region as well as elsewhere in the world. For these reasons, the potential and likely critical habitats for these species and others may be expected to change too. Over the medium term, identification of likely impacts and of mitigation options for development activities will require periodic reassessments to be undertaken based on latest information on species concentrations and habitat use preferences as well as environmental predictors.

## ACKNOWLEDGEMENTS

This research was carried out with the financial support of Royal Dutch Shell. The authors wish to thank Leon Bennun and John Pilgrim (Biodiversity Consultancy), and Szabolcs Nagy (Wetlands International) for their valuable and constructive inputs. This research would not have been possible without the International Wader Study Group (Arctic Birds Breeding Conditions Survey), Goose Specialist Group, Swan Specialist Group and data contributors, or reviewers, including: Georg Bangjord, Tom Barry, David Boertmann, Stuart Butchart; Preben Clausen, Peter Cranswick, Guttorm N. Christensen, Mindaugas Dagys, Sergey Dereliev; Tim Dodman, Garry Donaldson, Bart Ebbinge; Anthony David Fox,, Tony Fox, Gwen Fox, Peter Glazov, Larry Griffin, Gudmundur A. Gudmundsson, Ward Hagemeijer, Richard Hearn, Tombre Ingunn, Sergei Kharitonov, Tyler Kydd, Kees Koffijberg, Richard Lanctot, Elena Lebedeva-Hooft, Jesper Madsen, Nina Mikander, Gerard Muskens, Valentin Ilyashenko, David Boertmann, Anders Mosbech, Hans Meltofte, Leif Nilsson, Ib Krag Petersen, Nicky Petkov, Peter Prokosh, Jouke Prop, Diana Solovyeva, Kristinn H Skarphéðinsson, David Stroud, Evgeny Syroechkovsky, Tero Toivanen, Yvonne Verkuil, Doug Watkins, Hannah Wauchope, Martin Wikelski, and Ramunas Zydelis. In addition, we appreciate telemetry work included in our analyses that was originally undertaken with support from the Akvaplan-niva, The Arctic Centre, Fund King Leopold III of Belgium for the Exploration and Conservation of Nature, Norwegian Polar Institute, Norwegian Institute for Nature Research (NINA), University of Groningen, The Governor of Svalbard, The British Schools Exploring Society, and The Norwegian Biodiversity Information Centre.

## DATA ACCESSIBILITY

1. MaxEnt scripts and R scripts are included in Appendix S1.
2. MaxEnt model (5x).
3. GeoTIFF rasters probability (5x).
4. Shapefiles

a. Likely Critical Habitat
b. Potential Critical Habitat

## Appendix S1 MaxEnt script to extract random background points

D:\MaxEnt\EnvLayersFinal>java-cp D:\MaxEnt\maxent.jar density.tools.RandomSample 100000 bioclim01.asc bioclim02.asc bioclim03.asc bioclim04.asc bioclim05.asc bioclim06.asc bioclim07.asc bioclim08.asc bioclim09.asc bioclim10.asc bioclim11.asc bioclim12.asc bioclim13.asc bioclim14.asc bioclim15.asc bioclim16.asc bioclim17.asc bioclim18.asc bioclim19.asc dist2coast.asc dist2estuary.asc dist2freshwater.asc dist2mudflat.asc elevation.asc elevation_aspect.asc elevation_slope.asc elevation_std.asc globcover2009_v2.3.asc soilgwrb_freshw.asc > 100krandomsample.swd

## R commands for Spearman’s ranked correlations

Sample = read.csv(file.choose(), header=Τ)
coefficients = cor(sample, method = “spearman“)
write.csv(coefficients, file=“coefficients.csv“)

## Occurrence data sources

abbcs
AEWA report Yamalo-Nenetsky Autonomous Okrug
NOF
ZMO
telemetry
ArtDatabanken
BioFokus
BirdlifeFinland
CLO
DN
FMNH
iNaturalist
MFU
Miljϕfaglig Utredning
naturgucker
NRM
miljolare
gbif
Kolguev
Mindaugas Dagys

## Appendix S2

**Table S1.** *Overview of environmental predictors*

| Predictor code | Description | Reference |
| --- | --- | --- |
| *BioClimatic variables* |  | (Hijmans *et al.* 2005) |
| Bioclim01 | Annual Mean Temperature |  |
| Bioclim02 | Mean Diurnal Range (Mean monthly (max temp - min temp)) |  |
| Bioclim03 | Isothermality (BIO2/BIO7) (* 100) |  |
| Bioclim04 | Temperature Seasonality (standard deviation *100) |  |
| Bioclim05 | Max Temperature of Warmest Month |  |
| Bioclim06 | Min Temperature of Coldest Month |  |
| Bioclim07 | Temperature Annual Range (BIO5-BIO6) |  |
| Bioclim08 | Mean Temperature of Wettest Quarter |  |
| Bioclim09 | Mean Temperature of Driest Quarter |  |
| Bioclim10 | Mean Temperature of Warmest Quarter |  |
| Bioclim11 | Mean Temperature of Coldest Quarter |  |
| Bioclim12 | Annual Precipitation |  |
| Bioclim13 | Precipitation of Wettest Month |  |
| Bioclim14 | Precipitation of Driest Month |  |
| Bioclim15 | Precipitation Seasonality (Coefficient of Variation) |  |
| Bioclim16 | Precipitation of Wettest Quarter |  |
| Bioclim17 | Precipitation of Driest Quarter |  |
| Bioclim18 | Precipitation of Warmest Quarter |  |
| Bioclim19 | Precipitation of Coldest Quarter |  |
| *Euclidian distances* |  | Derived |
| Dist2coast | Distance to nearest coast |  |
| Dist2estuary | Distance to nearest estuary |  |
| Dist2freshwater | Distance to nearest freshwater body |  |
| Dist2shallowflatcoast | Distance to shallow coastal flats |  |
| *Elevation and elevation-derived* |  |  |
| Elevation | Elevation from Bioclimatic variable | (Hijmans *et al.* 2005) |
| Elevation_slope | Elevation slope | Derived |
| Elevation_std | Elevation standard deviation | (Danielson & Gesch 2011) |
| *Others* |  |  |
| GlobCover2009_v2.3 | Global Landcover | (Arino *et al.* 2012) |
| Soilgwrb_freshw | Dominant soiltype | Derived |

*Table S2. Pairwise correlations of environmental layers*

## Appendix S3 Supplementary figures

**Fig. S1.**
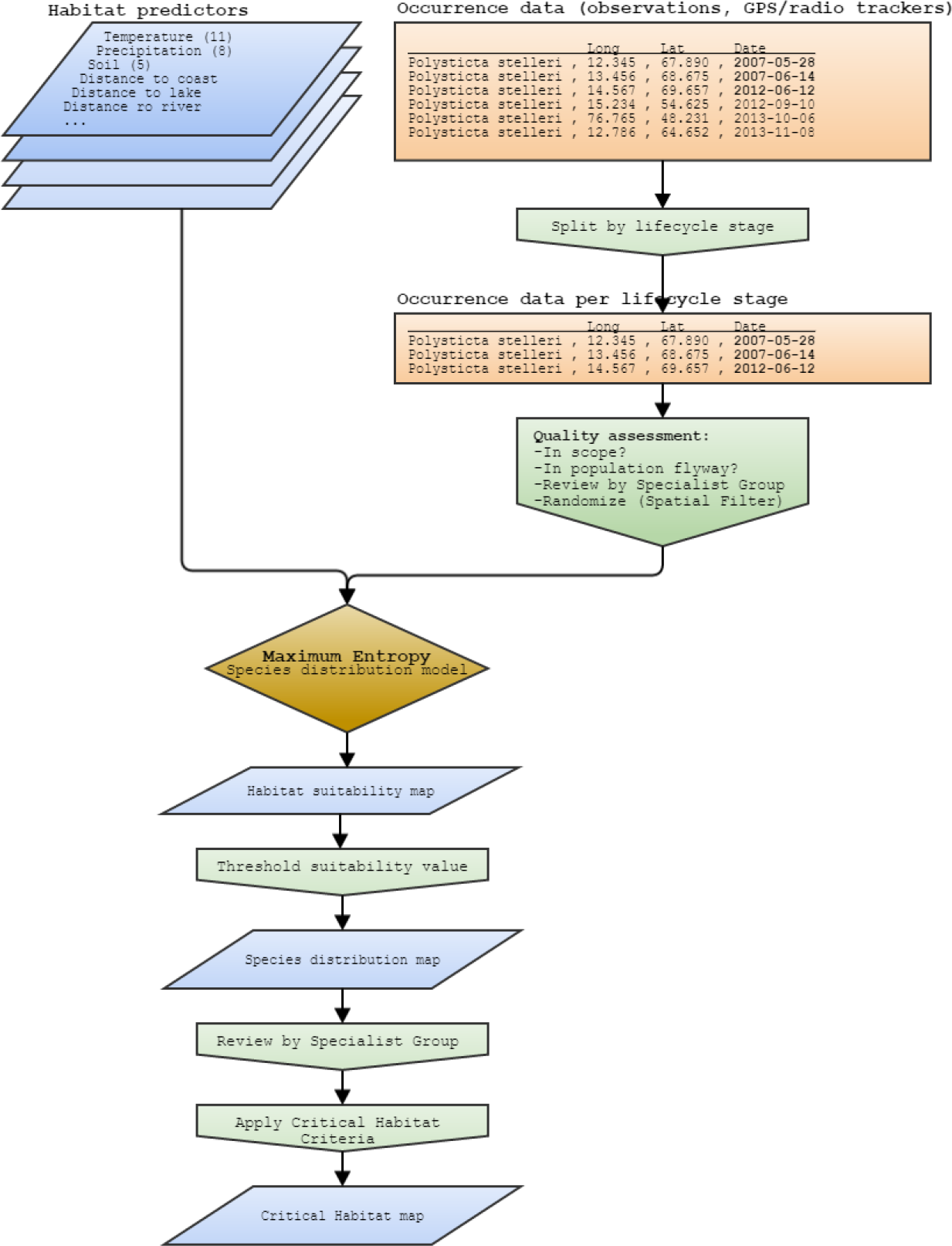
*Overview of the method used to create the critical habitat maps*

